# Investigating the Structural Effects of Anti-Thrombin Anticoagulant Aptamers on Activation of Human Prothrombin

**DOI:** 10.1101/2025.01.24.634711

**Authors:** Romualdo Troisi, Alessandro Cangiano, Nathan Cowieson, Vera Spiridonova, Pompea Del Vecchio, Luigi Paduano, Filomena Sica

## Abstract

Disorders of the blood coagulation remain a leading cause of death and disability worldwide raising the search for therapeutic agents able to modulate the coagulation cascade. Different oligonucleotide aptamers have been selected against different coagulation factors and some of them are in preclinical or clinical studies. In particular, anti-thrombin aptamers are promising drugs as they inhibit the activity of the α-thrombin and, simultaneously, limit thrombin production via prothrombinase by binding its precursor prothrombin. To investigate the interaction of these aptamers with prothrombin, we performed extensive analyses using calorimetric and spectroscopic techniques, which suggested that they recognize proexosite I of prothrombin and exosite I of thrombin with comparable affinity. SAXS experiments performed on the complex formed by the protein and NU172, the only anti-thrombin aptamer in advanced clinical trials, provided structural insights into aptamer-prothrombin recognition. Interestingly, the aptamer binding to proexosite I shifts the open-closed equilibrium of prothrombin toward the open conformation. A reasonable mechanism underlying the effects of anti-thrombin aptamers towards prothrombin conversion into thrombin has been proposed. Altogether, these results definitively qualify these aptamers as bitargeted drugs, being able to modulate both thrombin function and generation, and supply structural bases to design new anticoagulants, which lack health side effects.

## 1. Introduction

The coagulation cascade represents a sophisticated series of molecular events taking place in the blood in response to bleeding caused by tissue injury.^[1]^ The coagulation mechanism is regulated by several prothrombotic and antithrombotic events that assure the achievement of appropriate hemostasis.^[2–5]^ One key player in the regulation of hemostatic and non-hemostatic processes is thrombin (also known as factor IIa, FIIa), a multifunctional serine protease that performs its function by recognizing several interactors through the active site and two anion-binding sites (exosite I and II).^[6]^ This factor continues to be a key focus in cardiovascular therapy and a primary target for antithrombotic and anticoagulant treatments.^[7–13]^ Thrombin (**Figure 1**A) is produced in the common pathway of the coagulation cascade from the prothrombin zymogen through proteolytic cleavages catalyzed by the prothrombinase complex.^[1,6,14]^

**Figure 1.**
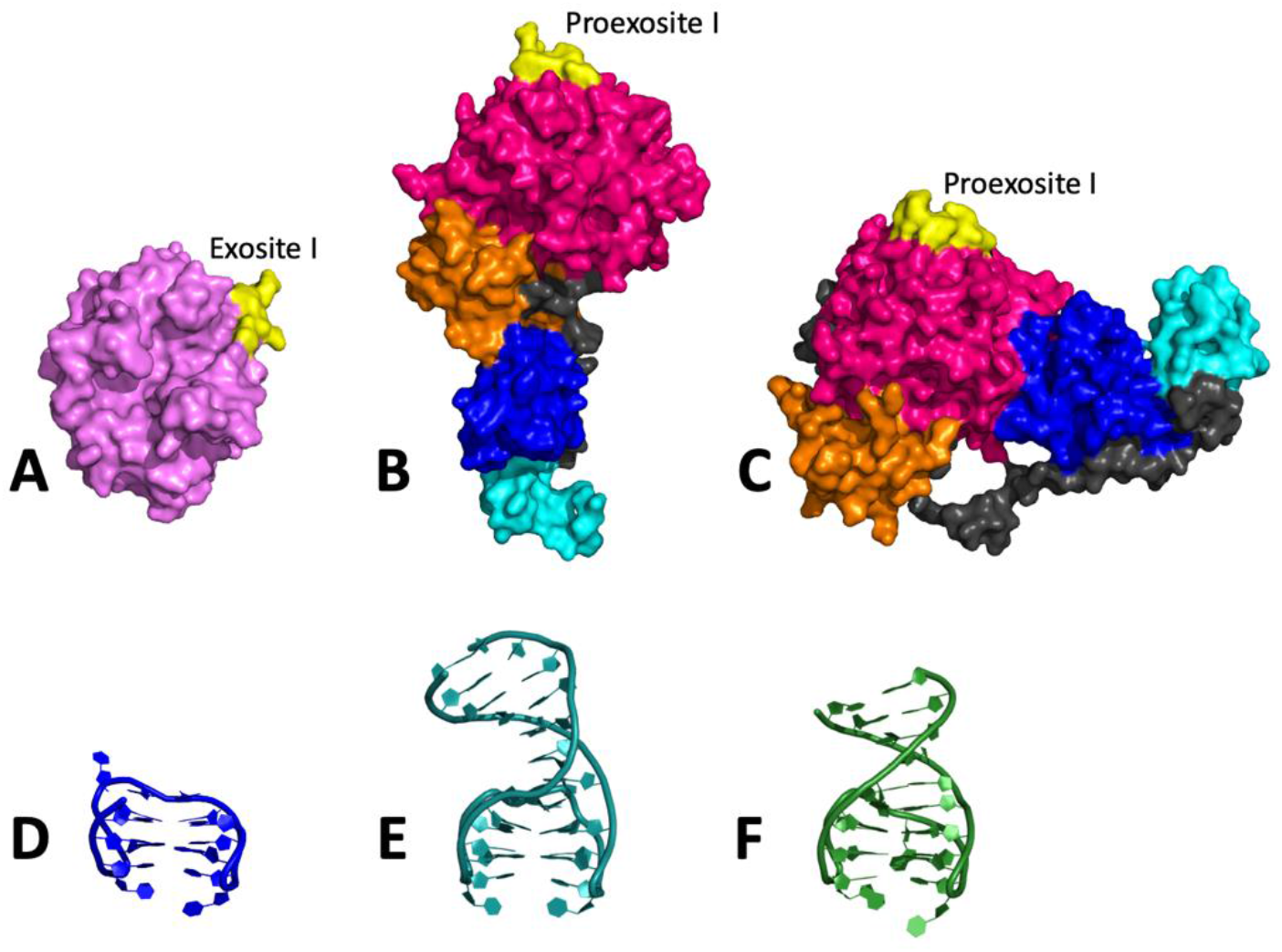
Surface and cartoon representation of A) thrombin (PDB code: 1PPB), prothrombin in the B) open (PDB code: 5EDM) and C) closed (PDB code: 6C2W) forms, D) TBA (PDB code: 4DII), E) RE31 (PDB code: 5CMX), and F) NU172 (PDB code: 6EVV). The GLA, K1, K2, and PD in prothrombin structures are cyan, blue, orange, and magenta, respectively, while exosite I of thrombin and proexosite I of prothrombin are yellow.

Prothrombin, or factor II (FII), is a modular protein organized into the membrane-anchoring γ-carboxyglutamic acid (GLA), the kringle-1 (K1), the kringle-2 (K2), and the protease (PD) domains (Figure 1B and Figure 1C), connected by three flexible linkers.^[15]^ A combination of mutagenesis, structural biology, and single molecule spectroscopy, has indicated that the linker 2, composed of 26 amino acids and connecting K1 to K2, strongly influences activation and conformational plasticity of prothrombin and enables the protein to sample alternative conformations.^[16–19]^ In particular, these studies have revealed that prothrombin exists in equilibrium between closed and open conformations. Specifically, in a L-shaped closed prothrombin (Figure 1C), K1 interacts with PD and closes the access to the catalytic pocket, whereas, in an I-shaped open form (Figure 1B), K1 is located far from PD, thus exposing the active site. The elongated open conformation is able to dimerize, to interact with specific membrane receptors through K1, and is more susceptible to proteolysis than the closed form, which is considered an autoinhibited form of the protein.^[15]^

The conversion of prothrombin to thrombin is induced by the interaction with prothrombinase, a complex composed of the enzyme factor Xa (FXa,) and the factor Va (FVa) assembled on the cellular surface in the presence of calcium ions.^[14]^ This conversion involves cleavage at two distinct sites, Arg271 and Arg320, along two alternative pathways that generate the zymogen precursor prethrombin-2 and the active enzyme meizothrombin, respectively.^[14,15,20]^ Interestingly, while the open conformation is preferentially cleaved at Arg271, the closed one is preferentially cleaved at Arg320.^[19]^ Although the two intermediates do not accumulate under physiological conditions,^[21]^ the dominance of either pathways influences the rate of thrombin generation.^[22–24]^ The prethrombin-2 mechanism is preferred on the surface of platelets,^[21]^ whereas the meizothrombin pathway prevails on red blood cells and endothelium.^[14,23]^

FXa is the trypsin-like proteinase of prothrombinase and catalyzes prothrombin conversion into thrombin. The membrane-dependent interaction of FXa with the FVa produces a nearly 10^5^-fold increase in the rate of thrombin formation at the physiological concentration of prothrombin.^[14]^ In the absence of FVa, activation proceeds preferentially along the prethrombin-2 pathway.^[25]^ A detailed structural picture of the initial encounter between prothrombin and prothrombinase along the meizothrombin pathway has been recently obtained by cryo-electron microscopy (cryo-EM).^[25,26]^

Maintenance of prothrombin as a zymogen requires the encryption not only of the active site but also of the two exosites to prevent the dysregulation of hemostasis.^[20]^ In prothrombin, while exosite II is occupied by K2, exosite I exists in a flexible state (proexosite I) that is ordered upon conversion to thrombin.^[20,27]^ Proexosite I is already able to specifically bind peptide ligands and protease activated receptors (PARs) with an affinity substantially lower compared to that of the mature exosite I.^[28]^

In thrombin, exosite I represents the target of many anticoagulant agents that inhibit thrombin-catalyzed fibrinogen cleavage.^[6]^ Among them, oligonucleotide aptamers, adopting peculiar three-dimensional structures,^[29,30]^ are particularly interesting because of their powerful and antidote-controllable activity and low immunogenicity.^[31]^ Interestingly, studies conducted by Kretz *et al*. demonstrated that the prototype of anti-thrombin DNA aptamers, the G-quadruplex TBA,^[32]^ is able to recognize proexosite I and attenuates prothrombin activation by prothrombinase by over 90%.^[33]^ The same ability was found in the case of some duplex/quadruplex anti-thrombin DNA aptamers as well as an RNA aptamer, named R9D-14T, and a bivalent DNA aptamer.^[34–37]^ Since these aptamers inhibit both activity and generation of thrombin, they represent effective dual-targeting therapy agents that could provide a decrease of therapeutic doses and of bleeding rates.^[37,38]^

Here we present a comprehensive calorimetric/spectroscopic characterization of the prothrombin interaction with three anti-thrombin DNA aptamers: the above mentioned TBA^[32]^ (Figure 1D) and the duplex/quadruplex RE31^[39]^ (Figure 1E) and NU172^[40]^ (Figure 1F) aptamers. On the basis of crystallographic studies, these oligonucleotides similarly contact with the same protruding residues of the thrombin exosite I, exploiting a pincer-like system formed by a couple of two-residue lateral loops of the G-quadruplex domain.^[30]^ The effect of the aptamer binding to proexosite I on the prothrombin conformational equilibrium has been analyzed by using as ligand NU172, the only aptamer to advance to phase II clinical trials (ClinicalTrials.gov Identifier: NCT00808964), and employing Small-Angle X-ray Scattering (SAXS) experiments. Such experiments revealed that the protein preferentially adopts an open conformation when complexed with the aptamer.

## 2. Results

### 2.1. Thermodynamic Analysis of the Aptamer-Prothrombin Interaction

The thermodynamic parameters of the binding between the prothrombin and the anti-thrombin aptamers (TBA, RE31, and NU172) were obtained by Isothermal Titration Calorimetry (ITC) experiments (**Figure 2**). For comparison, the binding of the same aptamers with thrombin was likewise investigated. Consistent with previous suggestions,^[32,39,40]^ the present results prove that the three aptamers form with thrombin a 1:1 complex. The same stoichiometry was observed in the case of the interaction with prothrombin. The thermodynamic parameters (**Table 1**) showed that the binding of these aptamers is characterized by a high affinity and specificity as revealed by the negative values of Gibbs energy changes and by the even more negative values of the enthalpy changes. For all the studied aptamers, the enthalpy contribution to the binding compensates for the loss of unfavorable entropy contribution. The binding constants K_b_ (Table 1) show a similar recognition ability for the three aptamers, which seem to have a slightly lower affinity towards prothrombin than thrombin. The differences in the binding thermodynamic parameters are subtle, revealing that, except for minor rearrangements, the investigated aptamers interact with prothrombin proexosite I in a manner consistent with their recognition of thrombin exosite I. It is important to underline that previous SPR analyses revealed NU172 to have a higher affinity than the other aptamers for both thrombin and prothrombin.^[41,42]^ This is consistent with X-ray structural data, which showed that the ligand-exosite I interface, which is very similar for the three aptamers, in NU172 is further stabilized by the interaction between Arg75 and Gua13, which belongs to the three-residue GTA loop placed on the opposite side respect to the two two-residue loops in the aptamer G-quadruplex domain.^[40]^ Conversely, the enthalpy values obtained from the here reported ITC experiments do not account for these variations. However, it must be underlined that the enthalpy values determined by ITC are the result of the combination of the noncovalent interactions at the aptamer-protein binding interface and hydration effects. Similarly, the net entropy values represent the sum of various contributions including the release of water/ions on the interaction surface and the changes in the conformational freedom, as well as the loss of rotational and translational degrees of freedom, of both the aptamer and protein upon complex formation. Consequently, in such complex systems, larger surface areas do not always are associated with more favorable thermodynamic parameters.^[43]^

**Figure 2.**
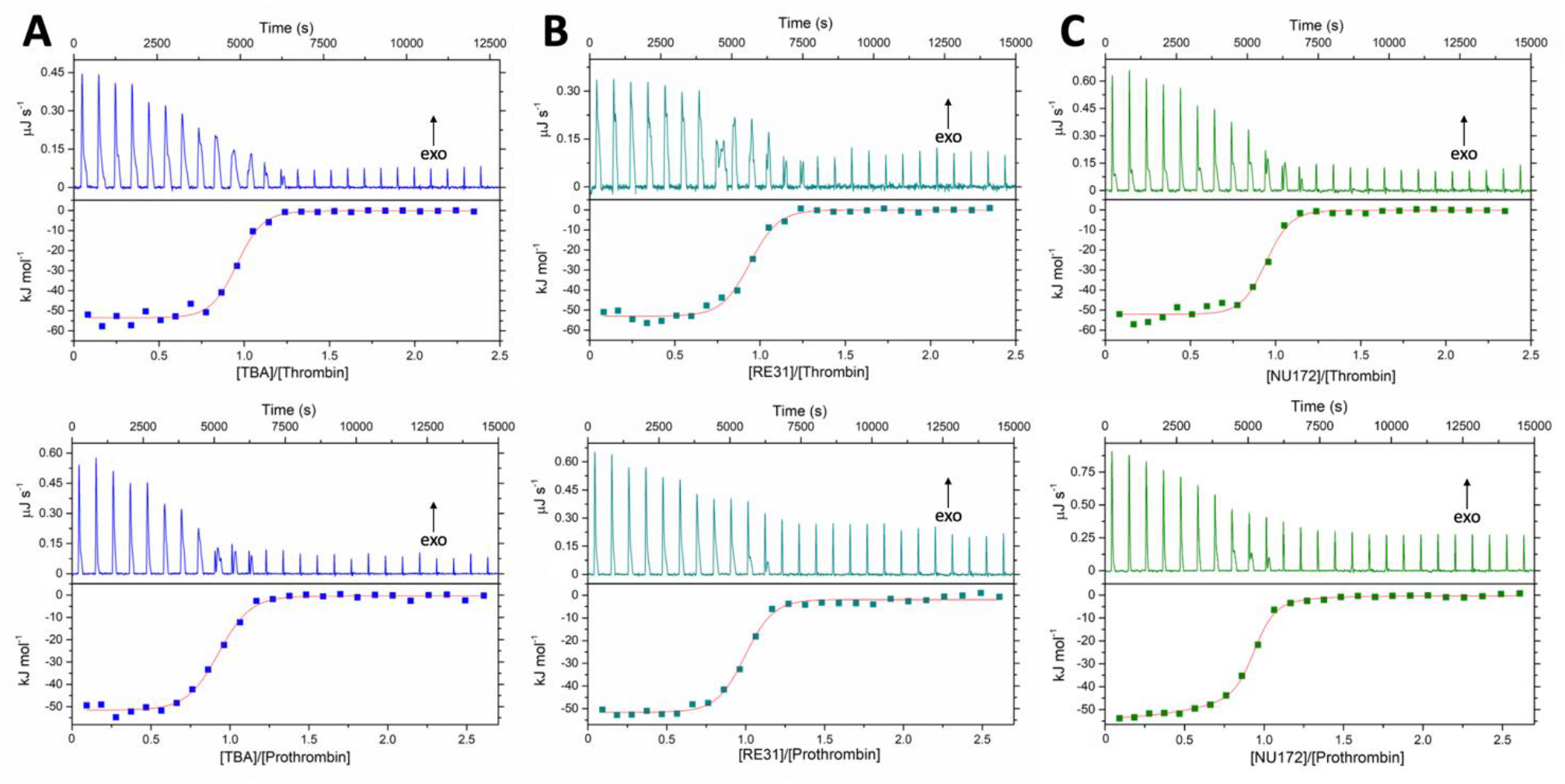
ITC binding isotherms for the interaction of A) TBA, B) RE31, and C) NU172 with thrombin (on the top) or prothrombin (on the bottom).

**Table 1.**
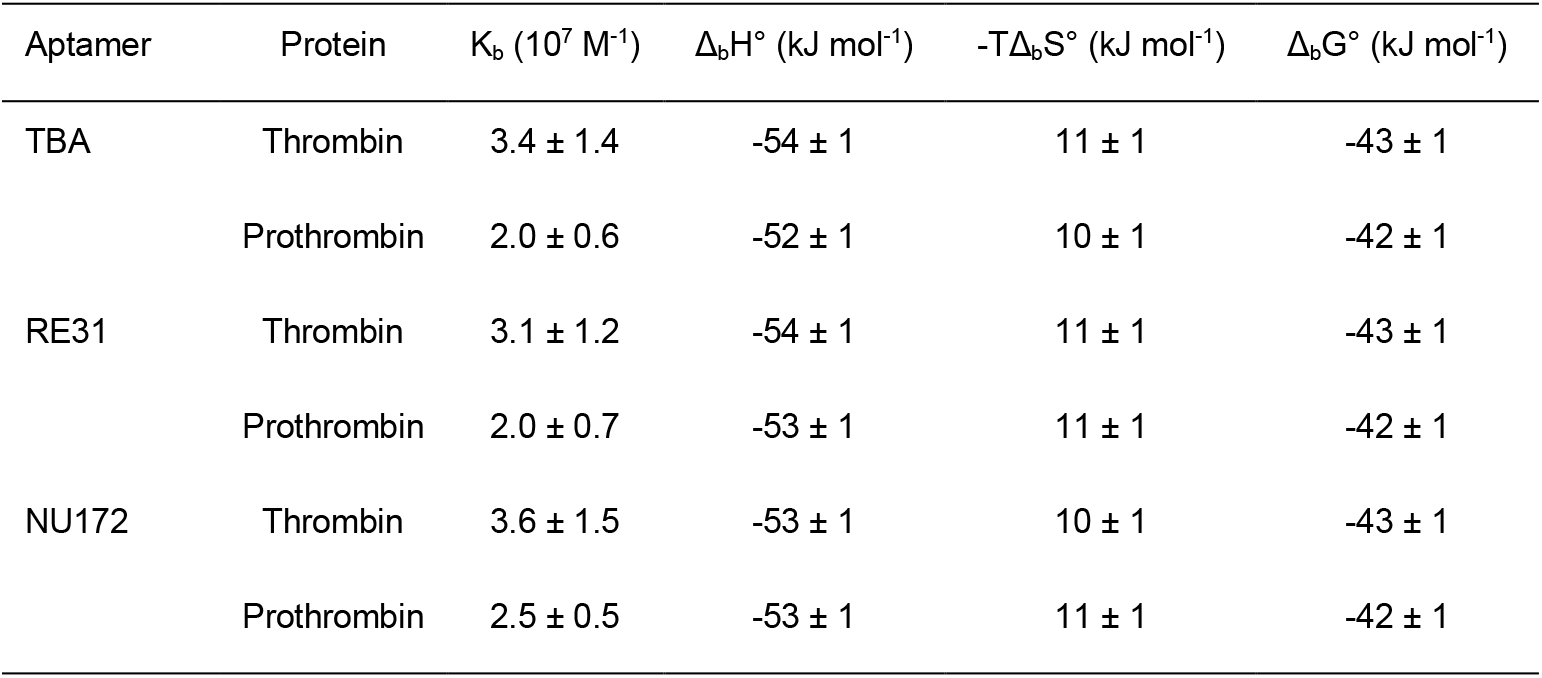
Thermodynamic parameters for the interaction of TBA, RE31, and NU172 with thrombin or prothrombin.

### 2.2. Effects of Prothrombin on Aptamer Folding

It has been widely reported in the literature that thrombin is able to act as a *chaperone* and to induce aptamer folding even in the absence of detectable amounts of cations, which stabilize the oligonucleotide architecture.^[44–46]^ To verify if this ability belongs also to prothrombin, Circular Dichroism (CD) spectra of TBA, RE31, and NU172 were recorded, in the absence of K^+^ or Na^+^ ions, before and after the addition of the protein. The spectra of the unbound TBA, RE31, and NU172 clearly indicate that in the absence of ions they do not adopt the G-quadruplex-based architectures observed in K^+^-rich solutions (Figure S1). Conversely, the presence of a positive signal around 290 nm, which is associated with the antiparallel G-quadruplex structure,^[47,48]^ characterizes the spectra of 1:2 aptamer-prothrombin solutions (**Figure 3**). Comparing these spectra with that of the free prothrombin (Figure S2), the signal around 290 nm appears only when the protein is bound to the aptamers. These spectral features (Figure 3) strongly suggest that the interaction of the prothrombin with the three aptamers promotes their correct folding as already observed for thrombin.^[44–46]^

**Figure 3.**
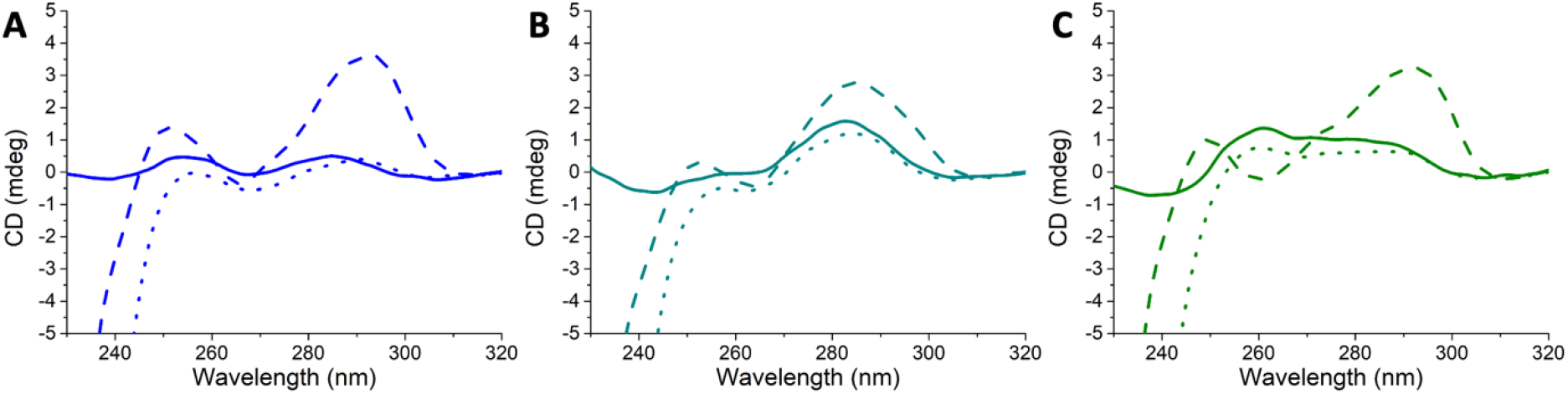
CD spectra of A) TBA, B) RE31, and C) NU172 in the absence (solid line) and in the presence of the native (dashed line) or denatured (dotted line) prothrombin. The spectra were recorded at 20 °C in 10 mM Tris-HCl pH 7.4, using an aptamer concentration of 1 μM. The prothrombin was added up to a concentration of 2 μM and denatured by heating the samples to 95 °C after the addition of 300 μM TCEP.

As a further control, the CD spectra (Figure 3) of these solutions were recorded after the denaturation of the protein, induced by heating the samples to 95 °C after the addition of 300 μM Tris(2-carboxyethyl)phosphine (TCEP), as reducing agent.^[49]^ The CD profiles clearly lose the antiparallel G-quadruplex signal observed upon aptamer binding with the native protein.

### 2.3. Effects of NU172 Aptamer on Prothrombin Conformation

Among the three examined aptamers, NU172 is certainly the most intriguing due to its high anticoagulant efficacy, which has prompted its consideration for clinical studies. Moreover, even if NU172 is an improved variant of TBA and RE31, the abovementioned ITC results and the indications of the crystallographic studies on aptamer-thrombin complexes^[32,39,40]^ suggest that the recognition motif between prothrombin and NU172 can be considered representative of that of the other two complexes. Starting from these considerations to obtain structural information about prothrombin and its complex with anti-exosite I aptamers in solution, the effects of NU172 binding to prothrombin on the enzyme conformational equilibrium were studied by SAXS analysis. This technique provides direct information about the size, shape, and oligomeric structure of biomolecules. The experimental data for intensity as a function of the modulus of scattering vector *q* were collected on the liganded and unliganded protein at three different concentrations (Figure S3). From the Fourier transformation of the scattering data, we calculated the pair distribution functions, P(r) (Figure S4), which give direct information about the molecule shape and size.^[50,51]^ The overall dimensional parameters for prothrombin and NU172-prothrombin complex are shown in Table S1. We observed a characteristic P(r) of an ellipsoid structure for both systems, showing a second low intensity peak that suggests the presence of a small percentage of aggregates (Figure S4). The envelopes of free and NU172-bound prothrombin were obtained from SAXS data and putative molecular models were built starting from the experimental crystal structures of free prothrombin in the open (PDB code: 5EDM) and closed (PDB code: 6C2W) conformations and of NU172-thrombin complex (PDB code: 6EVV). In this regard, it must be noticed that the high flexibility of the three linkers of prothrombin has prevented the single crystal X-ray diffraction analysis of the wild-type protein. However, a combined approach based on mutagenesis, functional studies, X-ray crystallography, and single-molecule FRET has allowed to identify the mutant proTΔ154-167 (PDB code: 5EDM), lacking 14 residues of linker 2, as prototype of the open form and the mutant proTS101C/A470C (PDB code: 6C2W), in which K1 is linked to the protease domain by an engineered disulfide bond, as prototype of the closed form.^[17–19]^ The final models of both the open and closed conformations of free prothrombin fit well the SAXS-derived envelope (**Figure 4**), suggesting a conformational heterogeneity of the protein in the solution.^[18]^ Notably, the fitting in the SAXS envelope of the prothrombin closed form produced a less compact conformation as compared to the relative crystallographic structure (Figure S5A). In contrast, after fitting, the complete model of open prothrombin retains an I-shaped structural organization in which the different domains tightly stack one on top of the other, similar to the PDB model (Figure S5B). The two NU172-prothrombin models were built by linking the aptamer on the prothrombin proexosite I in a similar orientation as found in the NU172-thrombin crystal structure. As shown in Figure 1B and Figure 1C, the relative position of proexosite I, which is part of PD, with respect to the other protein domains differs in the open and closed conformations. Differently from the model of the complex between NU172 and the closed prothrombin, the model involving the open form of prothrombin well fit in the extended envelope obtained by SAXS data (**Figure 5**), suggesting a preference of the aptamer towards the latter conformation. To further validate the aptamer binding to proexosite I and to exclude a different binding mode, we generated the model of the NU172-prothrombin complex using the recently released AlphaFold 3 (AF3) server.^[52]^ The program generated a model for the complex in which NU172 binds the closed form of prothrombin. In particular, the aptamer does not interact with proexosite I but with a PD region not too far from it (Figure S6). Based on this result, we shaped a new structural model of the complex by combining the open conformation of the protein refined by the SAXS data and NU172 at the binding site identified by AF3 program. Interestingly, neither the AF3-derived models (closed and open) of the NU172-prothrombin complex match the SAXS envelope (Figure S7), further corroborating our hypothesis that the aptamer interacts with prothrombin proexosite I as observed for thrombin exosite I.

**Figure 4.**
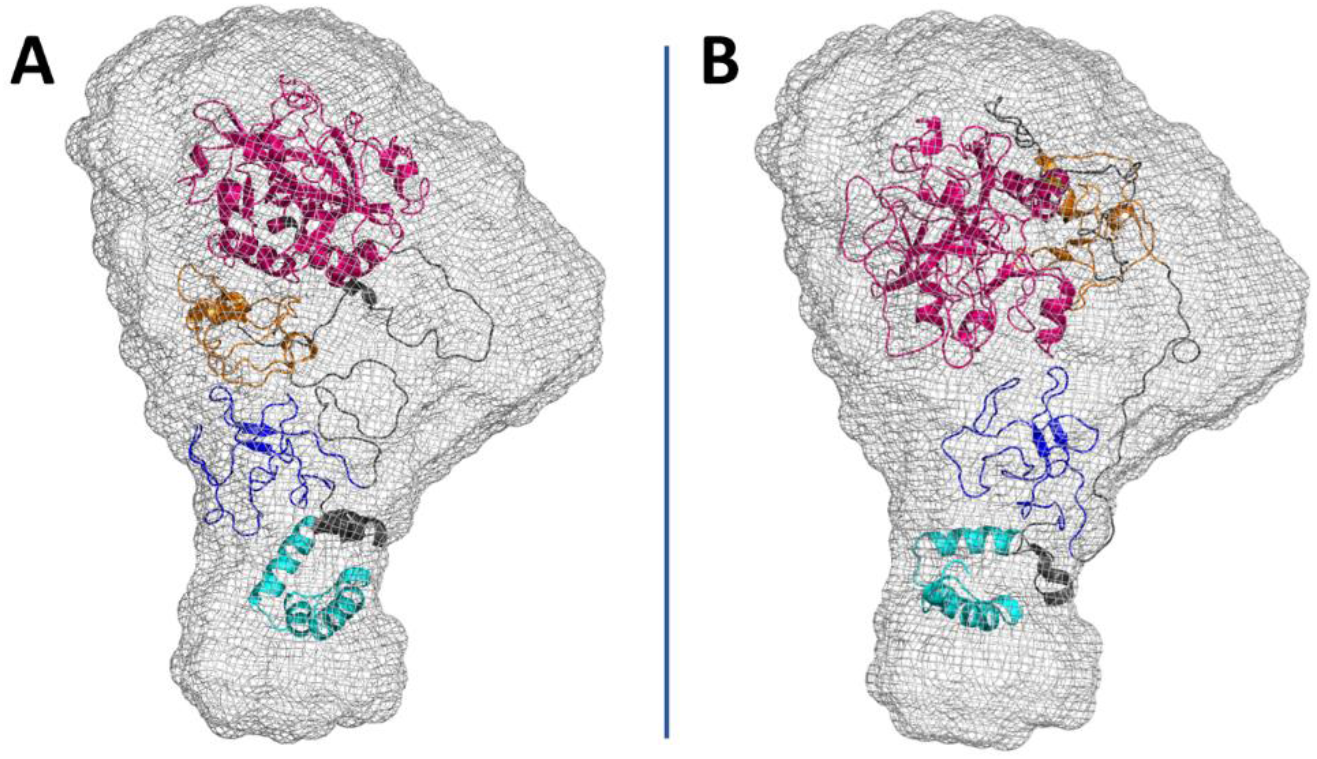
Reconstructed SAXS envelope of the free prothrombin superimposed on the atomic models of the protein in the A) open and B) closed conformations. The GLA, K1, K2, and PD are cyan, blue, orange, and magenta, respectively.

**Figure 5.**
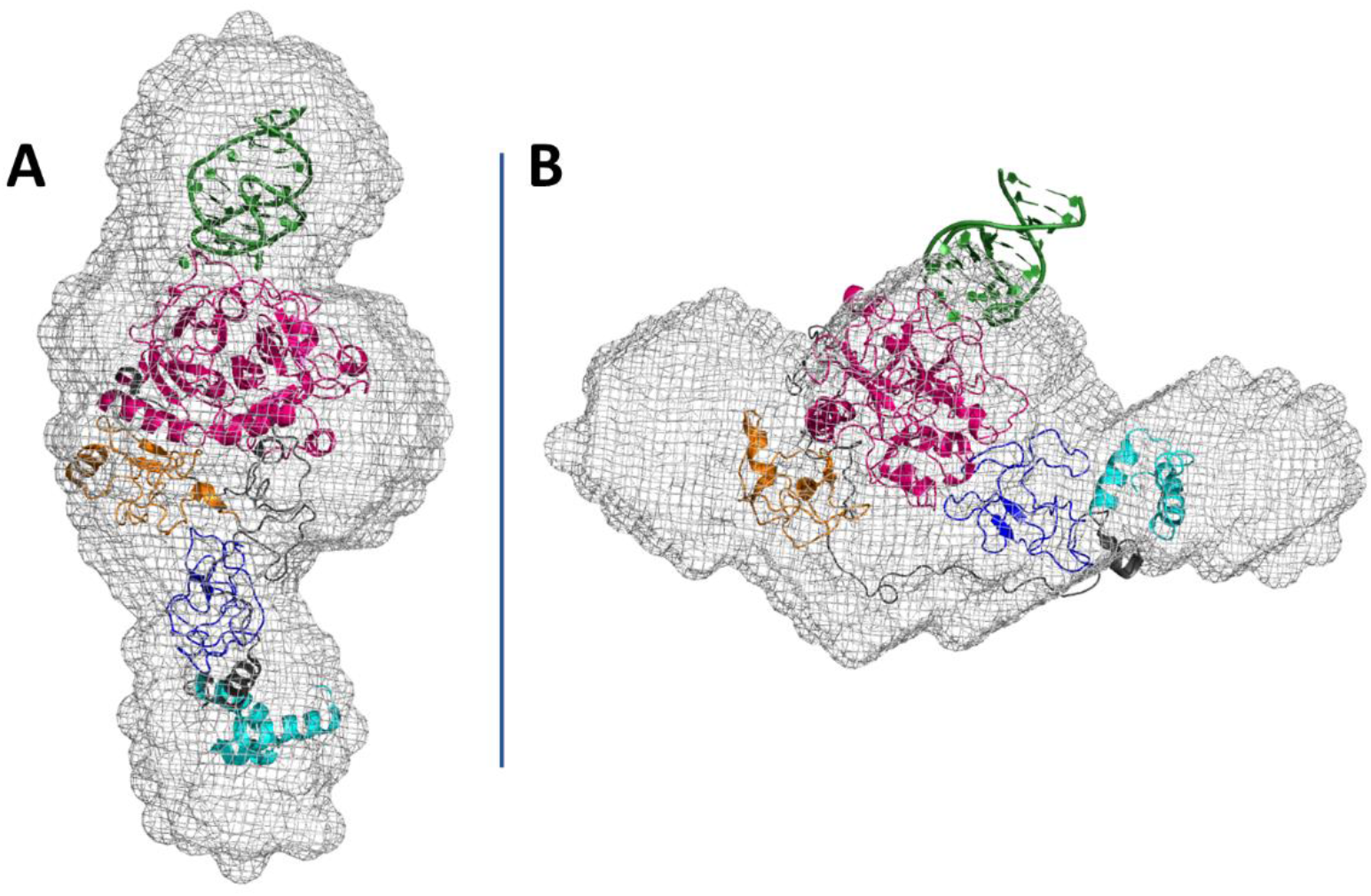
Reconstructed SAXS envelope of the NU172-prothrombin complex superimposed on the atomic models of the complexes formed by NU172 and the prothrombin in the A) open and B) closed conformations. NU172 is green, while the GLA, K1, K2, and PD of prothrombin are cyan, blue, orange, and magenta, respectively.

## 3. Discussion

Despite advances in anticoagulant therapy, disorders in the blood coagulation system remain the most common cause of death and disability in the world.^[53,54]^ Pathologic thrombosis or bleeding may occur whenever the hemostatic balance is disturbed due to various health conditions including surgery, trauma, malignancy, and congenital disorders.^[55–57]^ Anticoagulant drugs can inhibit thrombosis by altering various pathways within the coagulation system or by targeting thrombin directly, attenuating its function or generation.^[1,58,59]^ Current traditional anticoagulant therapy is often associated with bleeding complications, which must be quickly blocked by reversing the effect of the anticoagulant.^[1,60,61]^ In this context, aptamer–antidote system has received significant attention for its ability to control anticoagulation, and therefore bleeding, to the required degree.^[62–65]^

In this work, the results of ITC studies clearly indicate that some anti-thrombin anticoagulant aptamers can bind both thrombin exosite I and prothrombin proexosite I with comparable efficiency. CD experiments have also shown that prothrombin shares with thrombin a chaperone-like activity toward these aptamers. In addition, the recognition between anti-exosite I aptamers and prothrombin has also been characterized by SAXS. These experiments were carried out using NU172, the only anti-thrombin aptamer that has reached phase II of clinical trials.^[40]^ The results clearly show that, in the presence of the aptamer, the open-closed conformational equilibrium of prothrombin is shifted toward the open conformation. This phenomenon has been also observed in the presence of other ligands unrelated to aptamers. This is the case of argatroban, a thrombin-specific active site inhibitor.^[66,67]^ On binding to prothrombin, it shifts the conformational equilibrium towards the open form, the only one that has a catalytic pocket accessible to ligand binding.^[19]^ Argatroban also interferes with the interface interaction between K1 and PD that stabilizes the prothrombin closed form. Recently, it has also been found that the Fab domain of POmAb, a Type-I anti-prothrombin antibody,^[68]^ reduces thrombin generation by directly interacting with the K1 of the zymogen.^[69,70]^ Also, in this case, the binding hinders the K1/PD contacts and induces an increase in the relative abundance of the prothrombin open conformation. Interestingly, the binding of NU172 at the proexosite I region does not indicate close contacts in K1/PD interface region, nevertheless it does shift the conformational equilibrium of prothrombin towards the open form. A ligand-induced crosstalk between spatially distant regions such as exosite I/exosite II, exosite I/active site and exosite II/active site, has been well characterized in the case of thrombin and ascribed to the marked flexibility of the intervening bond connecting these regions within a thrombin-like skeletal organization.^[6,71–73]^ An analogous through-bond communication could occur when NU172 recognizes the proexosite I of prothrombin. Altogether, these data strongly suggest that even subtle modifications, induced by the ligand binding, can effectively modifies the relative stability of the two conformational forms. The aptamer-driven effect on prothrombin conformation revealed for NU172 can be confidently extended to TBA and RE31. Indeed, the here-reported thermodynamic data and the structural^[32,39,40]^ and dynamics^[71,72]^ studies described in the literature strongly suggest that the three aptamers could modulate the prothrombin conformation in a similar way.

By the here-presented results, interesting indications can be drawn regarding the way in which anti-thrombin aptamers are involved in the evolution of prothrombin into active thrombin. Although the prothrombinase complex, which controls the thrombin production, is not yet well characterized, a recent cryo-EM study has elucidated the structure of the complex formed by prothrombinase and the closed form of the thrombin zymogen.^[26]^ The authors were also able to derive a putative model for the binding of the prothrombin in the open form. In the closed form, the proexosite I is far away from FVa in apparent contrast with several studies in solution showing that FVa-proexosite I recognition has a controlling influence on the rate of prothrombin activation under physiological conditions.^[14,74–77]^ On the other hand, in the open form, proexosite I is in proximity of FVa leaving little space for additional ligands.^[26,78]^ Thus, it may be surmised that, along the prethrombin-2 pathway, the productive interaction between prothrombinase and the complex formed by the open form of prothrombin and an anti-exosite I aptamer, such as NU172, is disfavored. In addition, the aptamer-driven shift of the prothrombin equilibrium towards the open form, revealed by the SAXS analysis, may indirectly inhibit also the production of thrombin through the meizothrombin pathway.

Therefore, anti-thrombin aptamers are able to reduce the thrombin generation by acting simultaneously on the two transition pathways of prothrombin (**Figure 6**). Although this mechanistic model requires additional validations, it provides a consistent and global interpretation of the here-presented data, and the other structural and functional studies to date present in literature.

**Figure 6.**
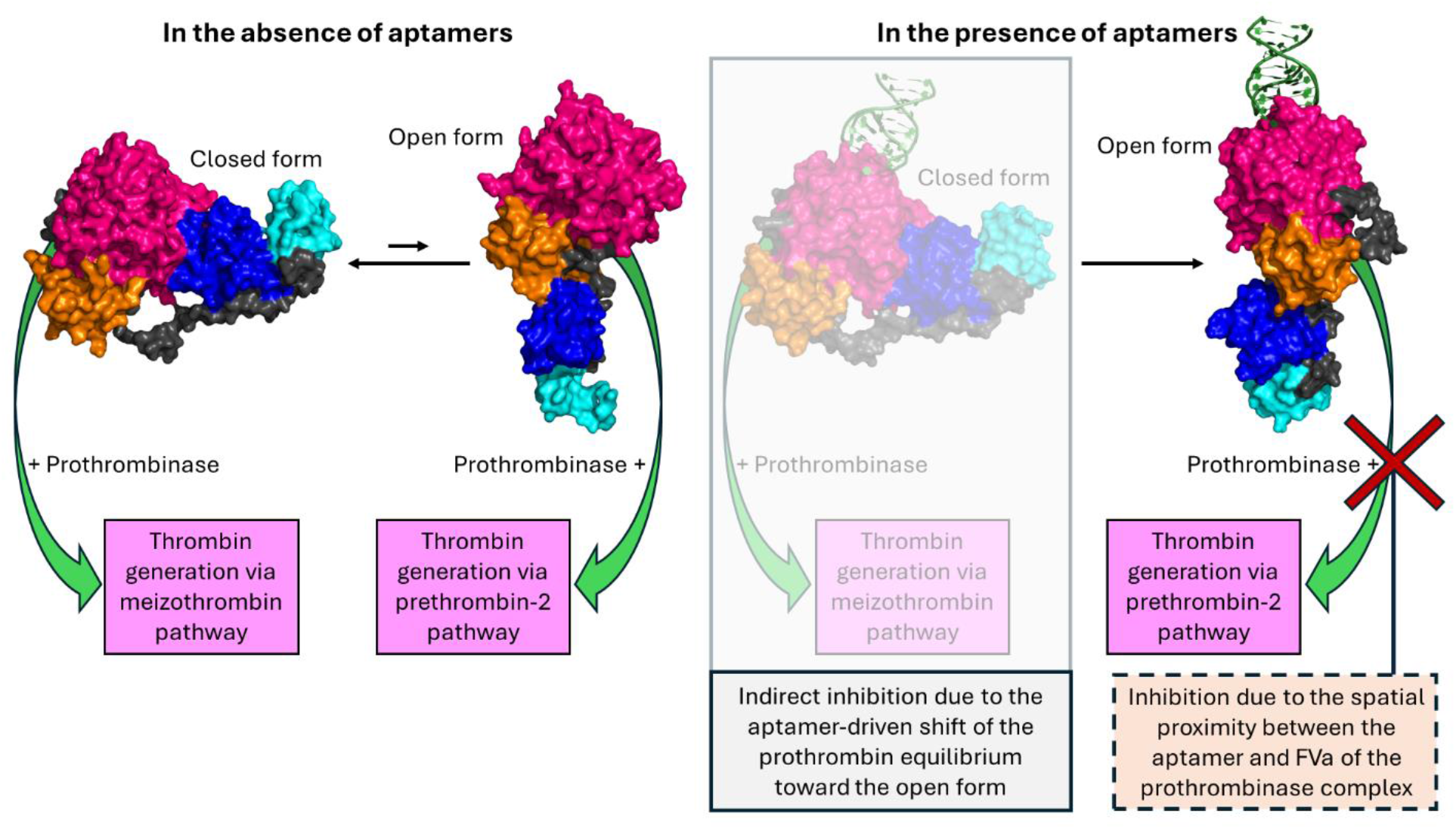
Schematic representation of the proposed mechanism of action for anti-thrombin aptamers in inhibiting thrombin generation. NU172 has a double action: stabilizes the protein open conformation and obstructs the correct assembly between this prothrombin form and prothrombinase. NU172 is green, while the GLA, K1, K2, and PD of prothrombin are cyan, blue, orange, and magenta, respectively.

These results definitively qualify anti-thrombin aptamers as bitargeted anticoagulants^[37]^ being able to efficiently inhibit thrombin generation and thrombin-fibrinogen recognition. Selective inhibition of thrombin formation by proexosite I binding aptamers would presumably have safer pharmacological profiles (i.e., less bleeding complications) compared to anticoagulants that inhibit thrombin activity directly.

In conclusion, this study has allowed to formulate reasonable structural hypotheses to explain the ability of thrombin anti-exosite I aptamers to modulate with high efficiency the prothrombin conversion into thrombin. In particular, we took advantage of the SAXS ability to offer structural information of biomolecules in solution, albeit at low resolution, to capture the effect of a ligand interaction on a protein conformation equilibrium that would be difficult to detect using other experimental approaches. These data can be fruitfully used for the design of a new generation of non-toxic and highly effective anticoagulants based on the use of the analyzed bitargeted aptamers, whose function could be easily blocked with the appropriate antidote.

## 4. Experimental Section

### Materials and Sample Preparation

The human prothrombin and the human D-Phe-Pro-Arg-chloromethylketone (PPACK)-inhibited thrombin were purchased from Haematologic Technologies (USA). TBA and NU172 aptamers were purchased from Merck (Sigma-Aldrich), while RE31 was purchased from Syntol (Russia). The concentration of protein and oligonucleotide samples were respectively determined by UV spectroscopy at 280 nm (at 20 °C) and 260 nm (at 90 °C), using molar extinction coefficients calculated from the primary sequences. Before each experiment the oligonucleotide samples were annealed by heating to 90 °C for 5 minutes, slowly cooling down, and then storing the samples at 20 °C overnight.

### Isothermal Titration Calorimetry

ITC measurements were performed at 25°C using a Nano-ITC III (TA instruments, New Castle, DE, USA) with a nominal cell volume of 961 µL. Protein (6.3 - 7.0 μM) and aptamer solutions (55 μM) were prepared in a buffer solution containing 100 mM KCl and 10 mM potassium phosphate at pH 7.4 and extensively dialyzed in the buffer solution at 4°C. The protein solution was loaded in the cell and titrated with the aptamer solution in the syringe by injecting 10 μL aliquots at 500 - 600 seconds intervals, for a total of 25 injections. The cell was stirred constantly at a rate of 250 rpm. The heat of dilution was measured in a separate experiment by adding the aptamer into the buffer solution under identical conditions. Raw data were integrated, corrected for non-specific heats, normalized for concentration, and analyzed by means of the NanoAnalyze software supplied by the manufacturer. Data analysis performed by means of an independent binding model gave the binding enthalpy (Δ_b_H°), equilibrium binding constant (K_b_), and binding stoichiometry (n). The Gibbs energy and the entropic contribution were then derived using the relationships Δ_b_G° = −RT ln K_b_ (R= 8.314 J mol^−1^ K^−1^, T = 298 K) and −TΔ_b_S° = Δ_b_G° − Δ_b_H°. ITC runs were repeated twice to evaluate the reproducibility of the results.

### Circular Dichroism

CD spectra were collected at 20 °C using a 1.0 cm path length cell on a Jasco J-715 spectropolarimeter equipped with a Peltier temperature control. Spectra were recorded with 50 nm min^-1^ scanning speed, 2 s response time, 1 nm data pitch, and 2.0 nm bandwidth. Each spectrum was obtained by averaging three scans.

### Small-Angle X-ray Scattering

For SAXS experiments, prothrombin and NU172-prothrombin samples were prepared in 10 mM Tris-HCl pH 7.4, 137 mM NaCl, 2.7 mM KCl, and 5 mM CaCl_2_, and concentrated to about 0.22, 0.44, and 0.88 mg mL^-1^ using 10 kDa-cutoff Centricon mini-concentrator (Vivaspin 500, Sartorius) and a refrigerated centrifuge (Microfuge 20R - Beckman Coulter). Following a standard protocol used for the preparation of aptamer-thrombin complexes,^[8,79,80]^ the NU172-prothrombin complex was prepared by placing a 2.5-fold molar excess of aptamer on a frozen sample of protein and leaving the sample at 4 °C overnight.

SAXS experiments were conducted at the B21 beamline of Diamond Light Source (Didcot, UK).^[81]^ The beamline was configured with a beam energy of 13.1 keV and a sample-to-detector distance of 3.7 meters, equipped with an EigerX 4M detector (Dectris). This setup enabled the collection of data for the scattering vector modulus *q=4πsin(θ/2)λ* ranging from 0.0045 to 0.34 Å^−1^, where *θ* is the scattering angle.^[81,82]^ SAXS data reduction was performed using the SCATTER IV software available at the B21 beamline. The radius of gyration (*R*_*g*_) was initially determined using the Guinier approximation for low-*q* (*qR*_*g*_<1.3) to assess data and sample quality.^[83]^ The pair correlation function *P*(*r*) was calculated using GNOM software,^[84]^ and the *R*_*g*_ values obtained from the *P*(*r*) analysis were compared with those from the Guinier analysis for further quality assessment.^[85]^ *Ab initio* shape reconstructions were performed using the DAMMIF program,^[86]^ employing dummy residues and constraints derived from the SAXS profile. For each dataset, a total of 20 reconstructions were generated and averaged using DAMAVER software to produce the most representative final model for each sample.^[87]^ The models were then refined with DAMMIN software.^[88]^

The modeling against the SAXS data was performed using AllosMod-FoXS^[89,90]^ and MultiFoXS^[91]^ programs and the Protein Data Bank (PDB) entries 5EDM^[17]^, 6C2W^[19]^, and 6EVV^[40]^, as sources for the starting models. In detail, prothrombin models (PDB codes: 5EDM, 6C2W) were manually curated and the complete models were obtained using the AllosMod-FoXS program. NU172-prothrombin starting models were built by superposing the catalytic domain of prothrombin from AllosMod-FoXS outputs and the thrombin heavy chain from NU172-thrombin crystal structure (PDB code: 6EVV). An energy minimization of the NU172-prothrombin complexes was performed using the YASARA Energy Minimization Server.^[92]^ Finally, a conformational sampling was carried out for all models using MultiFoXS program defining as flexible residues (47-64, 144-169, and 249-283) those belonging to the three intervening linkers of prothrombin.^[16]^

All structural figures were generated using PyMOL.

## Conflict of Interest

The authors declare no competing financial interests.

## Acknowledgements

A.C., N.S.C., and L.P. wish to thank Diamond Light Source, Didcot, Oxfordshire, England, United Kingdom for providing beamtime on B21 beamline (Reference SM-30763-2).

## Notes

### Competing Interest Statement

The authors have declared no competing interest.

