## Supporting Information for "Investigating the Structural Effects of Anti-Thrombin Anticoagulant Aptamers on Activation of Human Prothrombin"

[#]This work is dedicated to the memory of Vera Spiridonova, a co-author of this paper, who passed away in May 2023.

### Supplementary Table

**Table S1.** SAXS-derived dimensional parameters for prothrombin and NU172-prothrombin complex.

| Sample | P(r) analysis | | Guinier | Ellipsoidal fit | |
|---|---|---|---|---|---|
| | $R_g$ (Å) | $D_{max}$ (Å) | $R_g$ (Å) | Polar radius (Å) | Equatorial radius (Å) |
| Prothrombin (0.88 mg $mL^{-1}$) | 36.0±0.3 | 119.5 | 36.2±0.2 | 19.7±0.1 | 50.3±0.2 |
| Prothrombin (0.44 mg $mL^{-1}$) | 35.1±0.3 | 119.5 | 35.3±0.1 | 18.2±0.3 | 48.0±0.5 |
| Prothrombin (0.22 mg $mL^{-1}$) | 34.4±0.4 | 119.5 | 34.2±0.1 | 20.0±1.0 | 47.0±1.0 |
| Prothrombin+NU172 (0.88 mg $mL^{-1}$) | 41.3±0.4 | 138 | 41.0±0.2 | 11.1±0.2 | 53.5±0.1 |
| Prothrombin+NU172 (0.44 mg $mL^{-1}$) | 40.2±0.4 | 138 | 40.4±0.1 | 12.6±0.3 | 52.5±0.3 |
| Prothrombin+NU172 (0.22 mg $mL^{-1}$) | 39.0±0.5 | 138 | 39.1±0.1 | 13.0±1.0 | 53.3±0.6 |

$R_g$ = radius of gyration; $D_{max}$ = maximum dimension of the analyzed sample

### Supplementary Figures

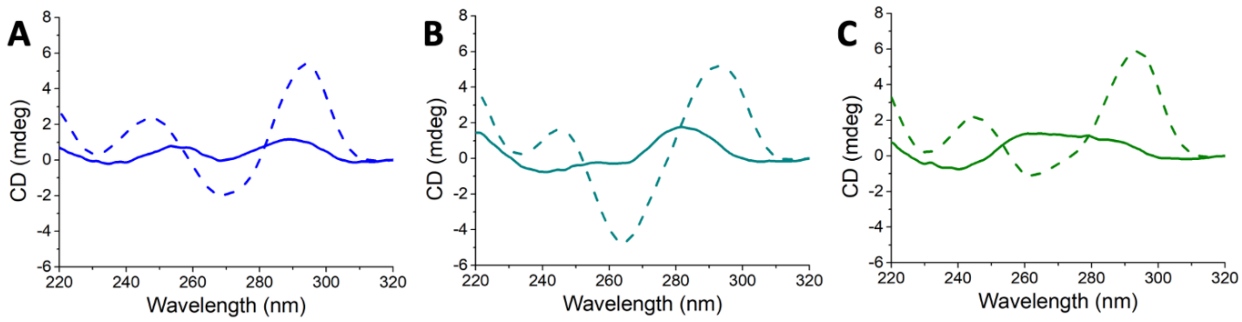


**Figure S1.** CD spectra of A) TBA, B) RE31, and C) NU172 in the absence (solid line) and in the presence (dashed line) of potassium ions. The spectra were recorded at 20 °C in 10 mM Tris-HCl pH 7.4 (solid line) or 10 mM potassium phosphate pH 7.4 and 100 mM KCl (dashed line), using an aptamer concentration of 1 μM.

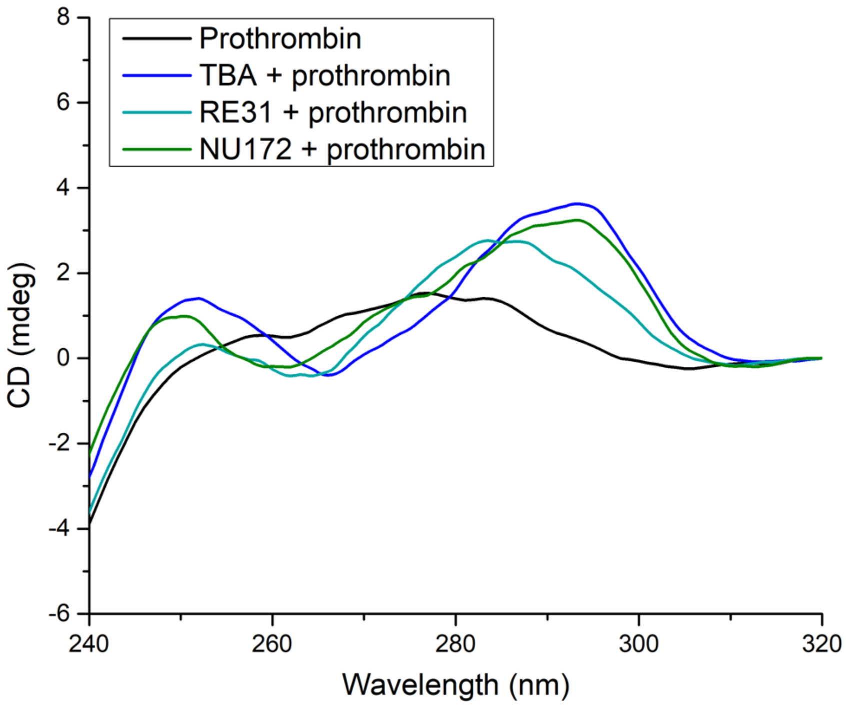


**Figure S2.** Comparison of CD spectra of free prothrombin and TBA, RE31, and NU172 in the presence of prothrombin. The spectra were recorded at 20 °C in 10 mM Tris-HCl pH 7.4, using an aptamer and protein concentration of 1 and 2 μM, respectively.

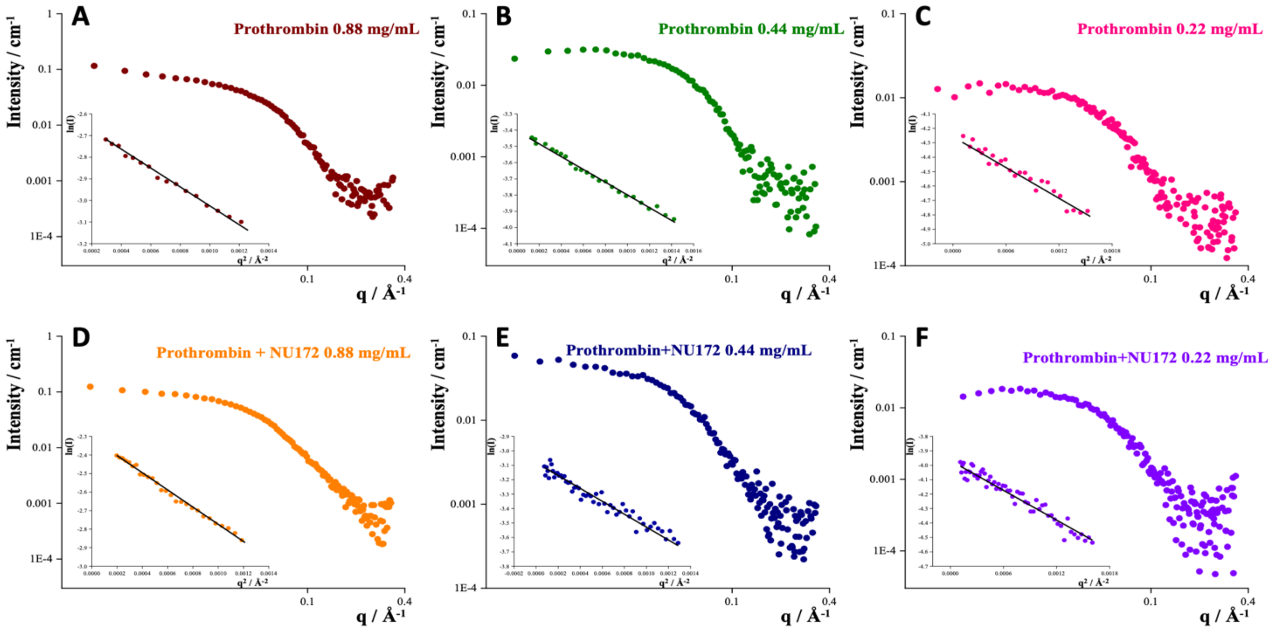


**Figure S3.** SAXS profiles of free prothrombin (A, B, and C) and NU172-prothrombin complex (D, E, and F). Guinier analysis of the low-*q* data is shown as insert in each graph.

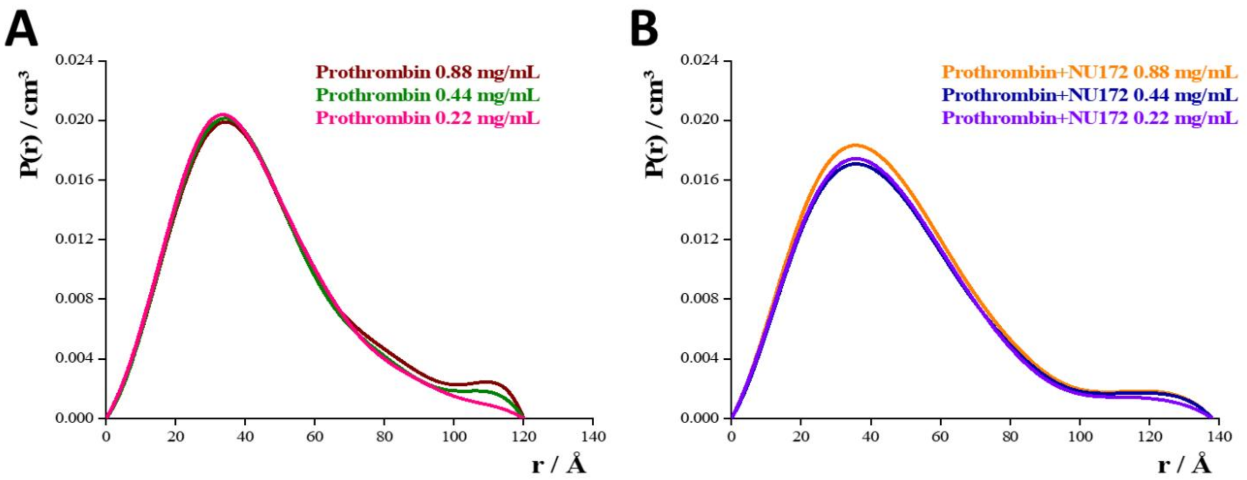


**Figure S4.** Pair distribution functions calculated from the scattering data for A) prothrombin and B) NU172-prothrombin complex.

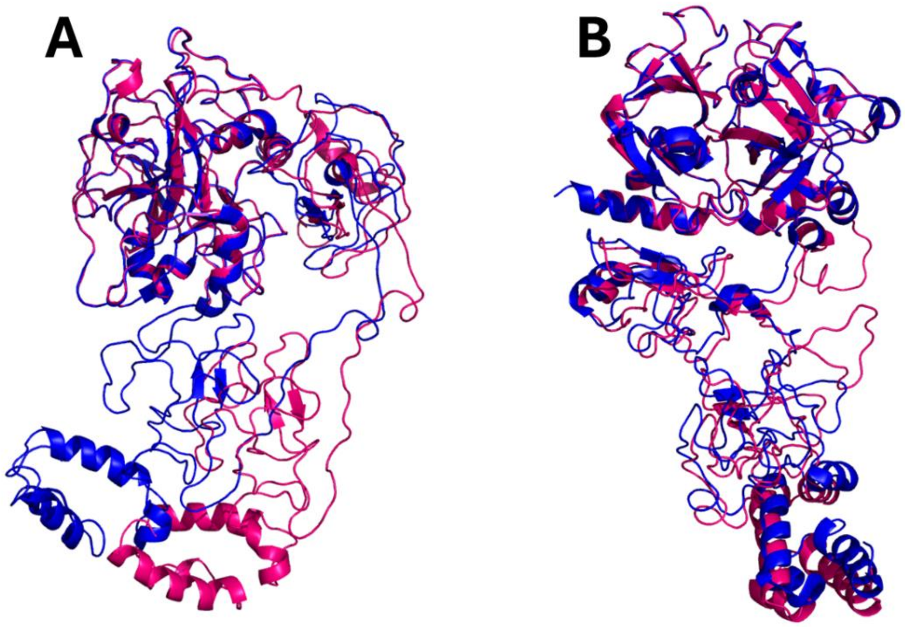


**Figure S5.** Superposition between the SAXS-derived models (magenta) and the crystal structures (blue) of prothrombin in A) closed (PDB code: 6C2W) and B) open (PDB code: 5EDM) conformations.

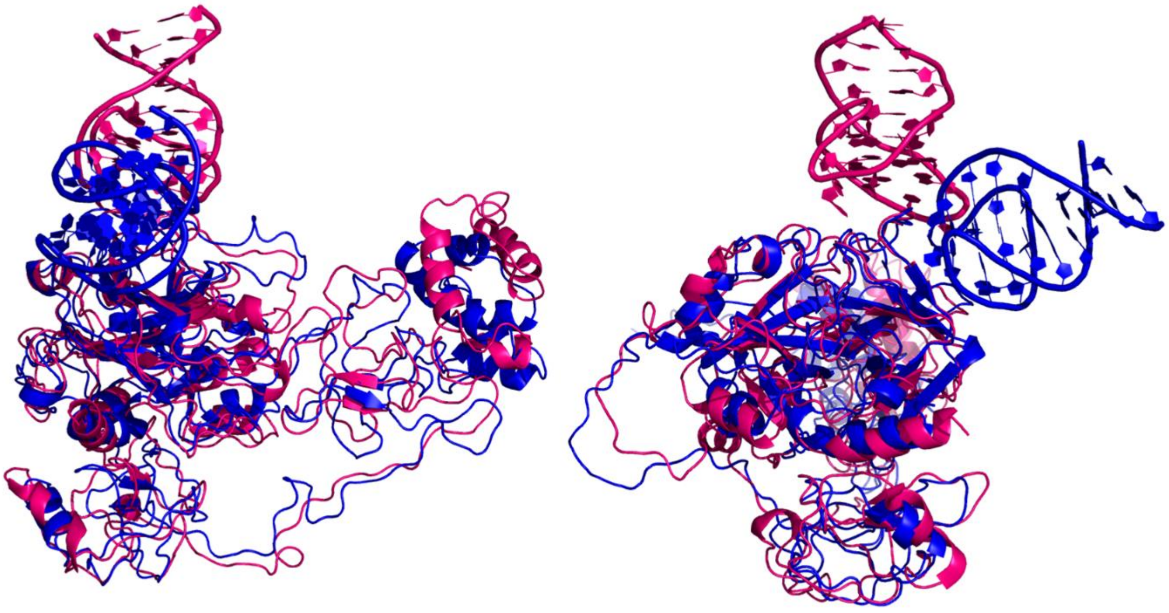

**Figure S6.** View in two orientations of the three-dimensional model of NU172-prothrombin complex predicted by AF3 (blue) superimposed on the SAXS-derived one (magenta). Prothrombin is in closed conformation. For the AF3 model, the predicted template modeling (pTM) and the interface predicted template modeling (ipTM) scores are 0.75 and 0.85, respectively.

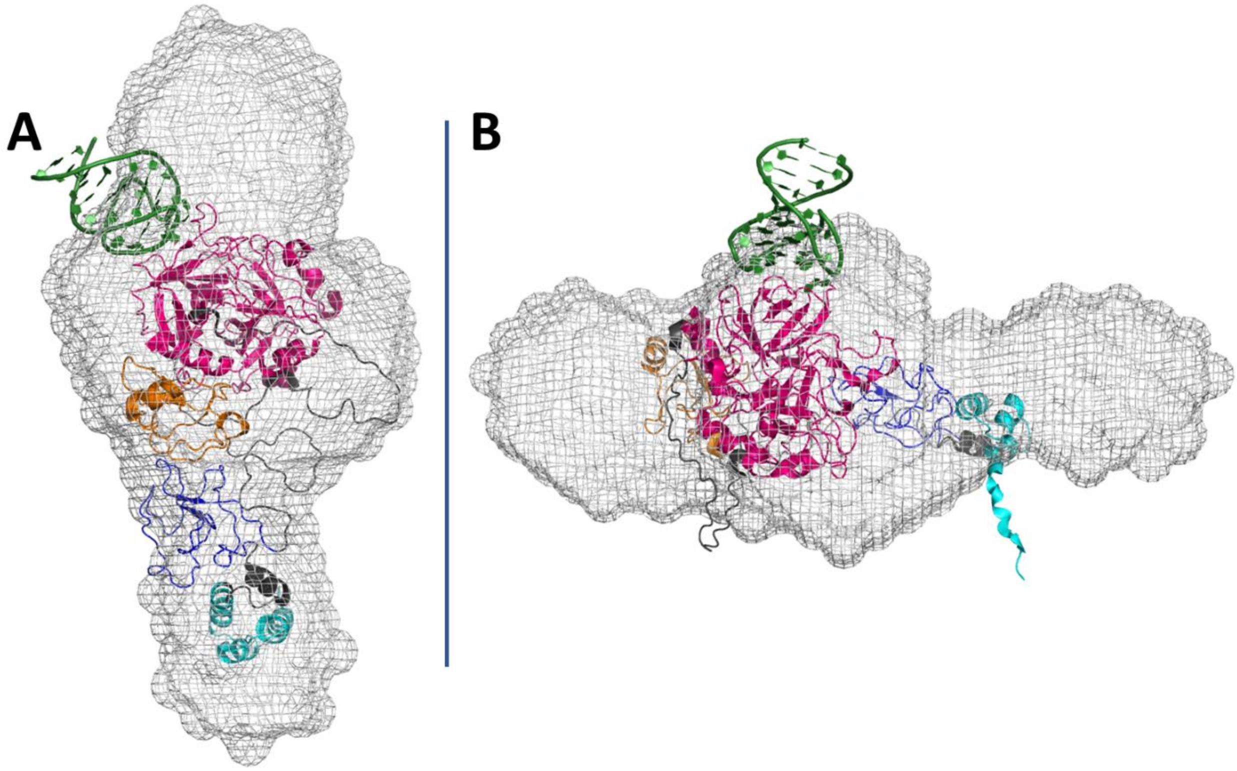


**Figure S7.** AF3-derived models of the complexes formed by NU172 and the prothrombin in the A) open and B) closed conformations superimposed on the reconstructed SAXS envelope of the NU172-prothrombin complex. NU172 is green, while the GLA, K1, K2, and PD of prothrombin are cyan, blue, orange, and magenta, respectively.
